# Predicting Antibody and ACE2 Affinity for SARS-CoV-2 BA.2.86 and JN.1 with *In Silico* Protein Modeling and Docking

**DOI:** 10.1101/2023.11.22.568364

**Authors:** Shirish Yasa, Sayal Guirales-Medrano, Denis Jacob Machado, Colby T. Ford, Daniel Janies

## Abstract

The emergence of SARS-CoV-2 lineages derived from Omicron, including BA.2.86 (nicknamed “Pirola”) and its relative, JN.1, has raised concerns about their potential impact on public and personal health due to numerous novel mutations. Despite this, predicting their implications based solely on mutation counts proves challenging. Empirical evidence of JN.1’s increased immune evasion capacity in relation to previous variants is mixed. To improve predictions beyond what is possible based solely on mutation counts, we conducted extensive *in silico* analyses on the binding affinity between the RBD of different SARS-CoV-2 variants (Wuhan-Hu-1, BA.1/B.1.1.529, BA.2, XBB.1.5, BA.2.86, and JN.1) and neutralizing antibodies from vaccinated or infected individuals, as well as the human angiotensin-converting enzyme 2 (ACE2) receptor. We observed no statistically significant difference in binding affinity between BA.2.86 or JN.1 and other variants. Therefore, we conclude that the new SARS-CoV-2 variants have no pronounced immune escape or infection capacity compared to previous variants. However, minor reductions in binding affinity for both the antibodies and ACE2 were noted for JN.1. We discuss the implications of the *in silico* findings and highlight the need for modeling and docking studies to go above and beyond mutation and basic serological neutralization analysis. Future research in this area will benefit from increased structural analyses of memory B-cell derived antibodies and should emphasize the importance of choosing appropriate samples for *in silico* studies to assess protection provided by vaccination and infection. Moreover, the fitness benefits of genomic variation outside of the RBD of BA.2.86 and JN.1 need to be investigated. This research contributes to understanding the BA.2.86 and JN.1 variants’ potential impact on public health. Taken together, this work introduces a paradigm for functional genomic epidemiology in ongoing efforts to combat the evolving SARS-CoV-2 pandemic and prepare for other hazards.

## Introduction

The continual emergence of SARS-CoV-2 variants remains a challenge for global public health initiatives. Notably, BA.2.86, which shares common ancestry with Omicron (BA.1), was initially identified in late July 2023 in Israel and Denmark (1, 2). Since then, it has disseminated globally (3–5). The initial worldwide spread of BA.2.86 sparked concerns regarding its potential impact on personal and public health. Acknowledging these concerns, the World Health Organization (WHO) designated BA.2.86 as a variant under monitoring on August 17th, 2023 (5).

The genetic makeup of BA.2.86 sets it apart from the original Omicron variant (B.1.1.529) and the XBB.1.5 variant (5), raising concerns due to its numerous mutations. These concerns are fueled by the observation of the rapid rise of previous variants with a high mutation count, exemplified by the surge in percent case counts of Omicron in late 2021 and early 2022, and XBB.1.5 in early 2023 (6, 7). BA.2.86 is distinguished by 35 mutations from the XBB.1.5 variant (8). Notably, the Spike gene (S) of BA.2.86 exhibits a distinctive profile with 33 mutations in comparison to the original Omicron variant, with 14 of these mutations occurring in the receptor binding domain (RBD) (8). Since antibodies primarily target the viral S protein’s RBD (9), it is considered a region of particular interest in the viral genome. Moreover, mutations in this vital RBD region can affect the efficiency of crucial viral-cell binding events (10), impacting the virus’ ability to bind to the human angiotensin-converting enzyme 2 (ACE2) receptor. For instance, the emergence of novel SARS-CoV-2 variants featuring RBD mutations has been associated with an increased affinity for binding to the ACE2 receptor in XBB.1.5 (10). This heightened RBD and ACE2 affinity has been linked to the accelerated person-to-person transmissibility observed in viruses like Omicron B.1.1.529 and XBB.1.5, leading to their predominance within the population (11). RBD mutations can also facilitate cross-species infections and increase the virus’ zoonotic potential (12). Furthermore, RBD mutations can hinder the efficacy of vaccines that target that region (13, 14).

Despite original concerns about the number of mutations in BA.2.86, and in a departure from past experiences, this variant has exhibited a notably low prevalence in the USA as of February 2024 (15). This indicates that relying solely on mutation counts does not suffice to gauge the severity of a new SARS-CoV-2 variant. More recently, the JN.1 variant, a relative of BA.2.86, emerged in 2023, once again sparking a concern owing to its increased mutation count. JN.1 is characterized by the Leu455Ser mutation in the RBD, which some authors have said makes it “one of the most immunologically evasive variants.” (16). Preliminary evidence indicates that the Leu455Ser mutation diminishes the binding affinity of the ACE2 receptor with JN.1’s RBD, potentially hindering viral entry into host cells while enhancing JN.1’s capacity to evade humoral immunity (17). Furthermore, Leu455Ser may confer greater evasion of JN.1 against the antibodies raised by a monovalent vaccine (18). In addition to JN.1’s mutation within the RBD, it bears three additional distinct mutations lacking in BA.2.86 (19).

Despite preliminary indications that RBD mutations would make BA.2.86 or JN.1 more capable than its preceding variants of escaping current vaccines, treatments, or antibodies produced by natural infection, conclusive empirical evidence is lacking and may not be immediately available. Furthermore, recent studies indicate that these new variants may not present pronounced differences in immune escape or infective capabilities when compared to previous variants despite their accumulated mutations. For example, (20) attested that JN.1’s serum neutralization escape did not increase over previously circulating strains despite its stronger increase in worldwide circulation when compared to BA.2.86 (16).

We build upon *in silico* methodologies established in our previous work (7, 21) to conduct extensive comparative analysis of the binding affinity of JN.1, BA.2.86, and other previous SARS-COV-2 variant RBDs (Wuhan-Hu-1, BA.1/B.1.1.529, BA.2, and XBB.1.5) to the human ACE2 receptor and neutralizing antibodies from infected individuals and patients vaccinated with different vaccines (including bivalent vaccines). Our findings hold significance for public health and contribute new methods to address the dynamic challenges posed by the evolving viral variants.

## Methods

In short, we examined 31 different RBD structures (27 RBD structures were extracted from 26 PDB files. One RBD structure was taken from a previous publication. Three PDB structures were predicted from six different SARS-CoV-2 variant RBDs (Wuhan-Hu-1, BA.1/B.1.1.529, BA.2, XBB.1.5, BA.2.86, and JN.1). Whenever more than one RBD structure was available for a variant, a single representative was selected based the presence of the interfacing residues and the results of docking validation experiments against the corresponding antibody. We cluster together BA.1 and B.1.1.529 in this study as they have identical RBD sequences. *In silico* docking experiments were used to calculate the binding affinity metrics between the six selected RBD structures and 27 different ligands, including 17 neutralizing antibody structures and ten ACE2 structures, resulting in 162 docking experiments total. All the analyzed structures are available on GitHub (see “Data Availability Statement”). The complete methodology is detailed below.

### Viral Proteins

Given the high infectiousness of the Omicron subvariants and their predominance in the past two years, we selected BA.1/B.1.1.529, BA.2, XBB.1.5, BA.2.86, and JN.1 variants’ RBDs as well as the original Wuhan-Hu-1 strain (referred to here as “wild type” or WT) for docking. Table 1 summarizes the sources of different RBD structures, as detailed below.

**Table 1.**
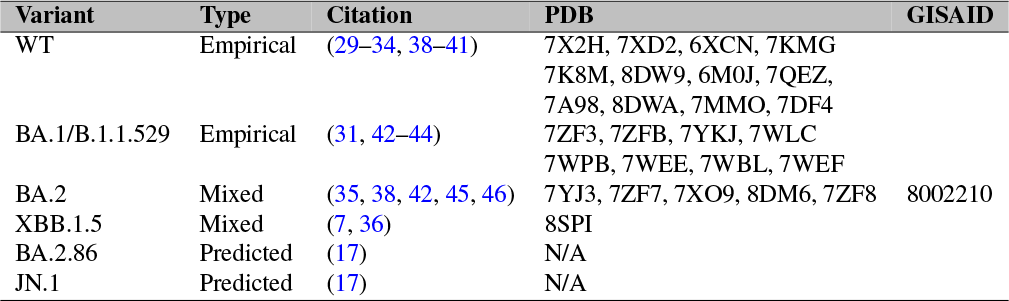
Viral RBD structures. WT indicates “wild type,” or Wuhan. Whenever more than one RBD structure was available for a variant, a single representative was selected based on sequence length, interfacing motif completeness, and the results of docking validation experiments against the corresponding antibody. The six RBD structures selected to represent each variant are available on GitHub (see “Data Availability Statement”).

We retrieved a complete genome sequence of BA.2 from the GISAID’s EpiCoV database (22). We annotated and translated the S gene following a similar method to Jacob Machado et al. 2021. We used the B.1.1.529 RBD sequence from our group’s initial *in silico* antibody docking study as a template, following the methodology described in (21). Finally, we extracted a fragment of the corresponding RBD region (within amino acid residues 336 through 528 in relation to the Wuhan-Hu-1 reference) (24). We obtained the RBD sequences for JN.1 and BA.2.86 from Yang et al. 2023. These sequences were used for structural protein prediction using AlphaFold2 (25) via ColabFold using default parameters (26). We relaxed the side chains in the ColabFold generated structure with the Amber relaxation procedure for docking (27).

We obtained the XBB.1.5 structure from our group’s previous *in silico* paper. The XBB.1.5 structure was generated using ColabFold (7). We downloaded other available SARS-CoV-2 Spike RBD crystal structures from the Protein Data Bank (PDB) (28). We derived the WT and BA.1/B.1.1.529 RBD structures from the Protein Data Bank as they represent empirically derived structures from an RBD-Antibody or RBD-ACE2 complex (29–38).

For every RBD-antibody and RBD-ACE2 complex we isolated on PDB, we extracted the RBD structure from that complex. We extracted the RBD structure from each complex and docked that structure against the corresponding Antibody or ACE2 structure to reproduce the initial complex for validation and ensure an accurate docking location.

### Antibody Selection

We expanded the antibody selection compared to previous studies that focused on therapeutic antibodies (7, 21).In this study, we selected 17 antibodies from the PDB, including those that were derived from vaccinated patients, patients vaccinated with breakthrough SARS-CoV-2 infection, and patients who experienced SARS-CoV-2 infection without vaccination. We included two therapeutic antibodies. 13 of the 17 selected antibodies were derived from memory B-cells from human patients (30, 31, 37, 42, 47). The initial complexed RBD was docked to each antibody structure alongside the representative RBD structures we derived from PDB and ColabFold. Antibodies are listed in Table 2.

**Table 2.** Selected antibodies from different conditions and their PDB structures. Note that the BNT162b2-CorV vaccine is commonly known as the “Pfizer-BioNtech” vaccine. Sinopharm developed BBIBP-CorV and was the first whole inactivated virus SARS-CoV-2 vaccination to obtain an emergency use authorization by the World Health Organization (30). The entries without a vaccine are antibodies obtained from unvaccinated patients before mass SARS-CoV-2 vaccination was available in the USA. The first 13 entries within the table are derived from memory B-cells collected from patients, whereas the last 4 entries were antibody structures generated from antibody sequences obtained from convalescent donors before mass vaccination or therapeutic antibodies developed for clinical use.

| Condition | Vaccine | Antibody | PDB | Citation |
| --- | --- | --- | --- | --- |
| Vaccinated | BBIBP-CorV | 6-2C | 7X2H | (30) |
| Vaccinated | BBIBP-CorV | 10-5B | 7XD2 | (30) |
| Vaccinated but infected with BA.1 | BNT162b2 | P3E6 | 7YKJ | (31) |
| Breakthrough infection |  |  |  |  |
| Vaccinated but infected with BA.1 | BNT162b2 | P2D9 | 8DW9 | (31) |
| Breakthrough infection |  |  |  |  |
| Vaccinated but infected with BA.1 | BNT162b2 | P1D9 | 8DWA | (31) |
| Breakthrough infection |  |  |  |  |
| Vaccinated but infected with BA.1 | BNT162b2 | Omi-3 | 7ZF3 | (42) |
| Breakthrough infection |  |  |  |  |
| Vaccinated but infected with BA.1 | BNT162b2 | Omi-31 | 7ZFB | (42) |
| Breakthrough infection |  |  |  |  |
| Vaccinated but infected with BA.1 | BNT162b2 | Omi-18 | 7ZFB | (42) |
| Breakthrough infection |  |  |  |  |
| Infected (strain not specified) | N/A | COVOX-150 | 7ZF8 | (42, 47) |
| Infected with WT | N/A | CV2.1169 | 7QEZ | (39) |
| Vaccinated | CoronaVac | XGv282 | 7WLC | (37) |
| Vaccinated | CoronaVac | XGv265 | 7WEE | (37) |
| Vaccinated | CoronaVac | XGv289 | 7WEF | (37) |
| Infected (strain not specified) | N/A | C105 | 6XCN | (33) |
| Infected (strain not specified) | N/A | C102 | 7K8M | (33) |
| Therapeutic | N/A | LY-COV1404 | 7MMO | (40) |
| Therapeutic | N/A | LY-COV555 | 7KMG | (41) |

### Human ACE2 Structures

We used PDB’s ACE2 structures derived from studies analyzing the structure of the ACE2-RBD complex. The ACE2 structures were isolated for docking. We docked the initial complexed RBD to each ACE2 structure alongside the RBDs that we derived from PDB and ColabFold. We selected ACE2 structures with the PDB IDs 6M0J, 7A^98, 7^DF^4, 7^YJ^3, 8^SPI, 7WBl, 7WPB, 8DM^6, 7^ZF7, and 7X09 (29, 34–36, 38, 42–46). We chose ten different structures of ACE2 to have an increased sample size with a diversity of minor structural deviations that may affect docking.

### Protein-to-Protein Docking

To prepare the Fab structures for docking, we renumbered the residues according to HAD-DOCK’s (v2.4) requirements such that there were no overlapping residue IDs between the heavy and light chains (48, 49). We selected residues in the Fab structures’ complementarity-determining regions (CDRs) as “active residues” for docking analysis to assess antibody neutralization. We selected residues in the ACE2 binding pocket forming polar contacts with the RBD in the crystallized structure as “active residues” for docking prediction and analysis of ACE2-RBD binding. We used the same active residues for each ACE2 and antibody structure for each dock to the six RBD structures. We selected residues in the S1 portion of the RBD as active residues. Each RBD has similar active residues when docking against an antibody. However, there are variations in active residue selection to account for differences in amino acid composition between variants. We docked each of the 17 antibody structures and 10 ACE2 structures against six RBD structures using HADDOCK, a biomolecular modeling software that provides docking predictions for provided protein structures (48, 49). Specifically, we used the HAD-DOCK Web Server (v2.4) for the docking simulations with the default parameters (48, 49).

The HADDOCK software produces multiple output PDB files of docking results and their subsequent docking metrics. We placed the top-scoring PDB output file, after qualitative analysis, to ensure a realistic binding pose for each docking experiment with HADDOCK into PRODIGY (v2.1.3) for further analysis. PRODIGY is a web service collection focused on binding affinity predictions for biological complexes (50, 51).

This process resulted in 162 sets of docked structures. We selected the top predicted output structure for each antibody-RBD or ACE2-RBD pair for quantitative analysis after assessing the binding pose. Statistical tests were conducted in R (52), implementing the Kruskall-Wallis and the paired Wilcoxon-Mann-Whitney test to compare different predictions (53, 54). We also used this top structure to visually analyze the structural conformation of interfacing residues and docked proteins using PyMol (v2.5.5) (55).

## Results

### Docking Results for assessment of Antibody and ACE2 binding Affinity

We compared docking predictions of viral proteins to antibodies and ACE2 with Kruskall-Wallis and paired Wilcoxon-Mann-Whitney tests. The tested variables were HADDOCK score, van der Waals energy, electrostatic energy, desolvation energy, buried surface area, and PRODIGY’s ΔG predictions. These tests return values that are not statistically significant between JN.1/BA.2.86 and previous Omicron variants at a 95% confidence level for every displayed metric, with a few exceptions, in Figures 1 and 2. The difference in Desolvation Energy is statistically significant, via the paired Wilcoxon-Mann-Whitney test, between XBB.1.5 and JN.1 in the antibody comparison, with JN.1 having a lower Desolvation Energy. The difference in desolvation energy between JN.1 and BA.1/B.1.1.529 is statistically significant in the ACE2 comparisons, with BA.1/B.1.1.529 having a lower desolvation energy. In the ACE2 analysis, the Kruskal-Wallis test yields statistically significant differences between all groups.

**Fig. 1.**
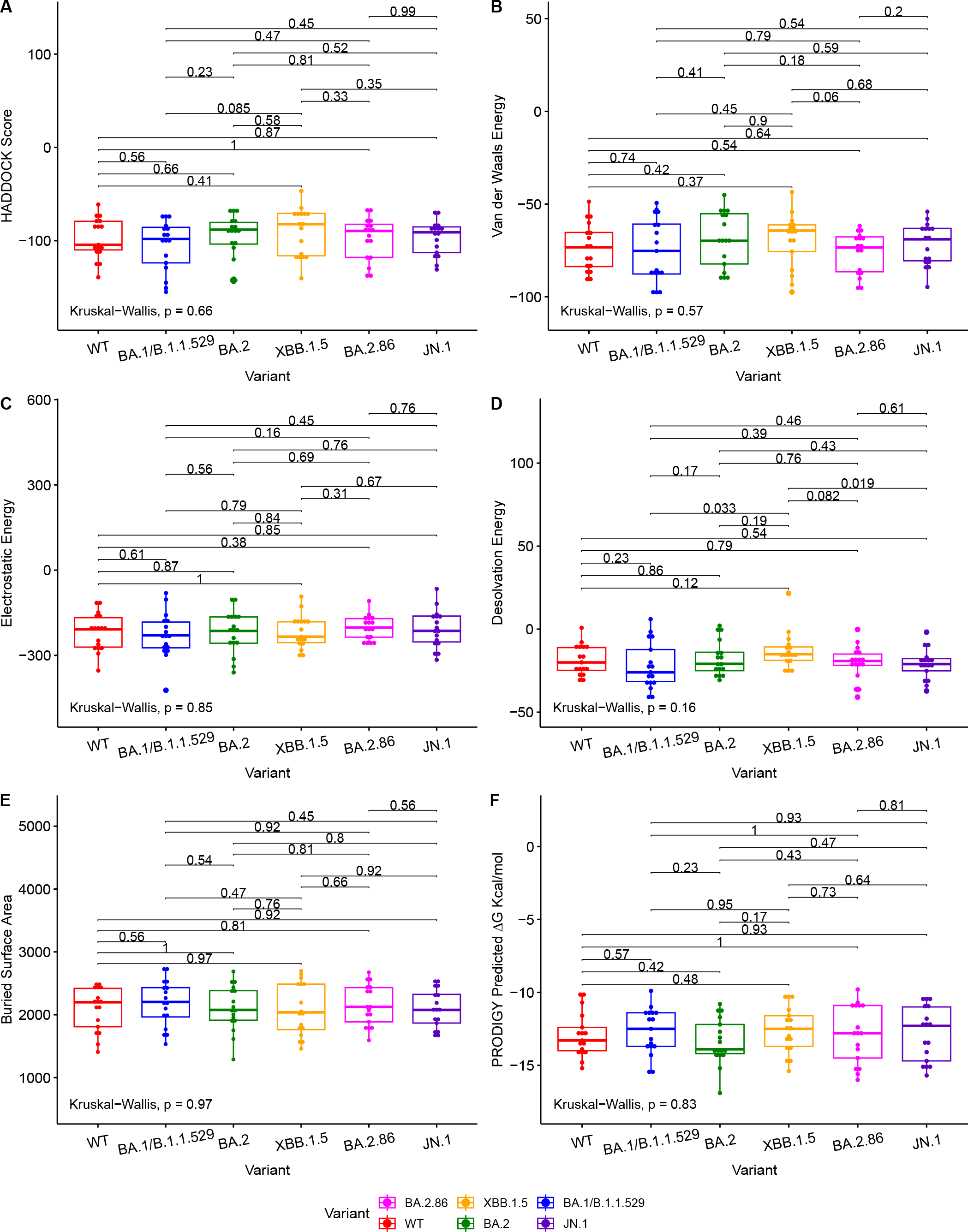
Boxplots to illustrate the comparative docking scores of antibody and RBD structures generated by HADDOCK and PRODIGY. Each boxplot highlights the distribution of docking scores for different variants. Pairwise comparisons were performed using the Wilcoxon-Mann-Whitney signed-rank test, indicated by the horizontal lines. The Kruskal-Wallis test was used to compare the independent samples’ medians, indicated in the bottom left.

**Fig. 2.**
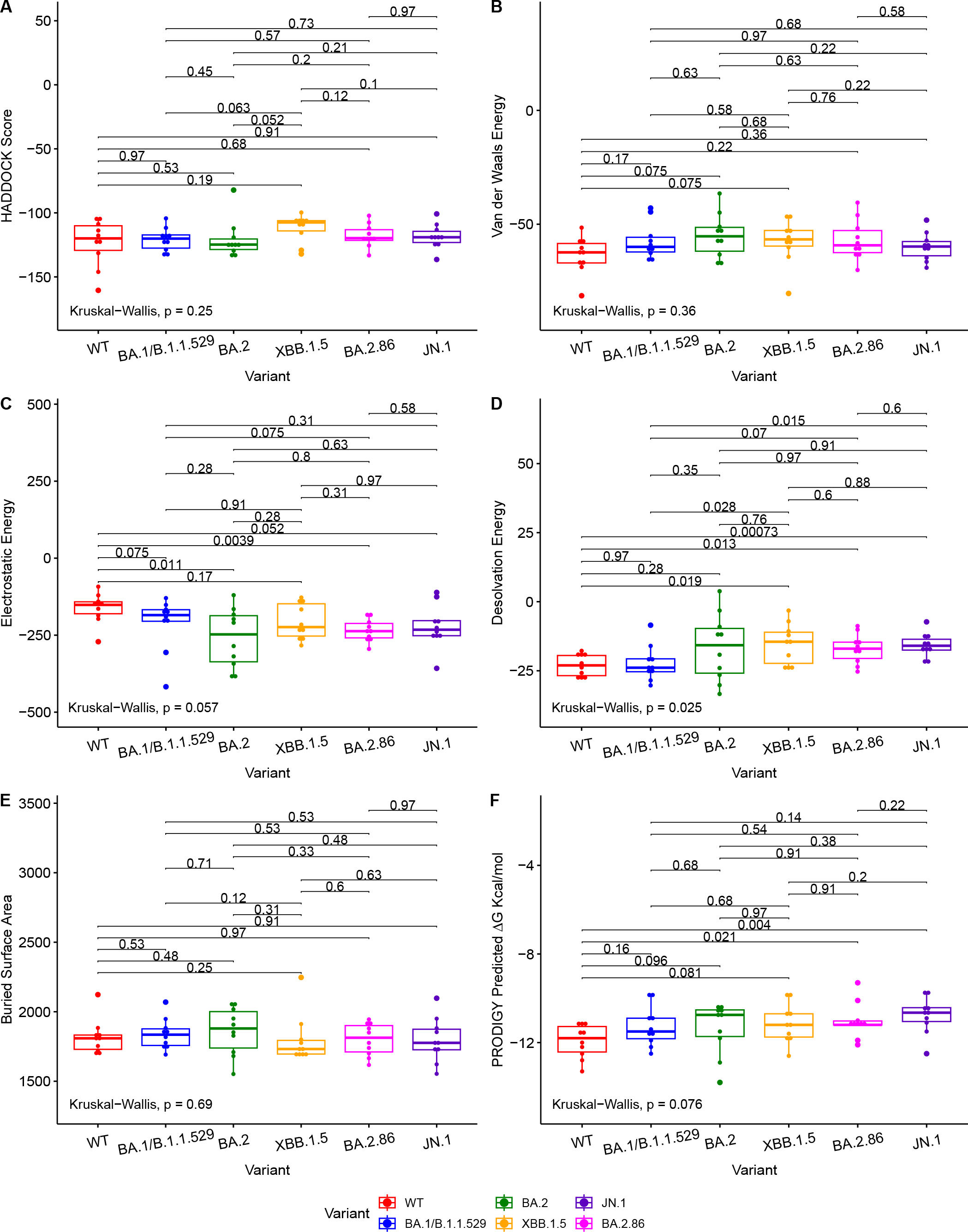
Boxplots illustrating the comparative docking scores of ACE2 and RBD structures generated by HADDOCK and PRODIGY. Each boxplot highlights the distribution of docking scores for different variants. Pairwise comparisons were performed using the Wilcoxon-Mann-Whitney signed-rank test, indicated by the horizontal lines. The Kruskal-Wallis test was used to compare the independent samples’ medians, indicated in the bottom left.

The Wuhan or WT variant exhibited a higher binding affinity, which is statistically significant, to ACE2 than the BA.2.86 and JN.1 variants when assessing ΔG value in Figure 2. The WT variant also exhibited a lower desolvation energy score when binding to ACE2, which is statistically significant, than the BA.2.86 and JN.1 variants. The WT variant exhibited a higher, statistically significant, electrostatic energy score than BA.2.86.

The Kruskal-Wallis test, which assesses statistical significance for group comparisons as a whole, returns a statistically significant *p* value at a 95% confidence interval in the ACE2 comparisons for Desolvation Energy in Figure 2.

We use the PRODIGY ΔG score as the primary metric to measure binding affinity. Thus, we conclude that all the Omicron subvariants’ (including BA.2.86 and JN.1) performance for the ACE2 and antibody docking simulations were similar. Slight reductions in antibody-RBD and ACE2-RBD binding affinity were observed for JN.1 when compared to BA.2.86, but they are not statistically significant at a p-value of 0.81 and 0.22, respectively, for a 95% confidence interval.

Figure 1 illustrates various metrics produced by HADDOCK and PRODIGY estimations of the protein-to-protein binding affinities between our aggregate antibody arsenal and RBD structures. It includes 17 antibody structures and six variant RBD structures (102 experiments in total). The results of ACE2 to RBD docking experiments are shown in Figure 2, including 10 ACE2 and six RBD structures (60 docking experiments in total). Figures 1 and 2 also show the non-significative *p*-values of the Kruskal-Wallis statistical test in the bottom left of each plot and the Wilcoxon-Mann-Whitney signed-rank test between each RBD, again with the exceptions mentioned above. These figures are derived from metrics obtained from the best PDB complex structure, determined by HADDOCK, for each experiment.

### Structural Analysis

Figure 3 shows the structural analysis of the interfacing residues between the RBDs of JN.1 and BA.2.86 with antibodies. The only difference in the RBD in JN.1 and BA.2.86 is the Leu455Ser substitution in JN.1. The residue at position 455 does not form polar contacts in either the JN.1-RBD or BA.2.86-RBD complexes in Figure 3 (A) or (B), nor is the residue at position 455 near the docking interface for either complex. The Leucine at position 455, in the RBD of BA.2.86, may interact with residues within the RBD. The substitution of Leucine for Serine at position 455 may alter intramolecular interactions of the RBD, inherently affecting its tertiary structure. This disparity in structure may affect the binding pose of the RBD in complex with the antibody, which is highly dependent on the antibody mechanism. In Figure 3(A), one can note that BA.2.86 generally forms polar contacts with residues that do not propagate steric hindrance. In contrast, Figure 3(B) shows that JN.1 forms two histidine polar contacts, a side chain that introduces more steric hindrance and rigidity. Histidine contacts may reduce proximal residue interdigitation, providing a lower binding affinity score. In Figure 3(C), the Phenylalanine at position 489 of BA.2.86’s RBD interdigitates well with antibody residues, potentially increasing nonpolar interactions and decreasing the effect of the steric hindrance from the bulky Phenylalanine side chain. In Figure 3(D), one can notice that residues appear less interdigitated than in Figure 3(C). This may be due to the destabilizing effect of the Serine at position 455 in JN.1, in addition to the alteration of the RBD tertiary structure mentioned previously. Leucine455 is shown in yellow in Figure 3(C), close to Isoleucine100 of the antibody. Both of these residues may form nonpolar interactions, strengthening binding. In 3(D), this Serine substitution has caused a lack of attraction between the two residues, potentially causing the segment within the RBD to retract back in docking and decrease binding affinity.

**Fig. 3.**
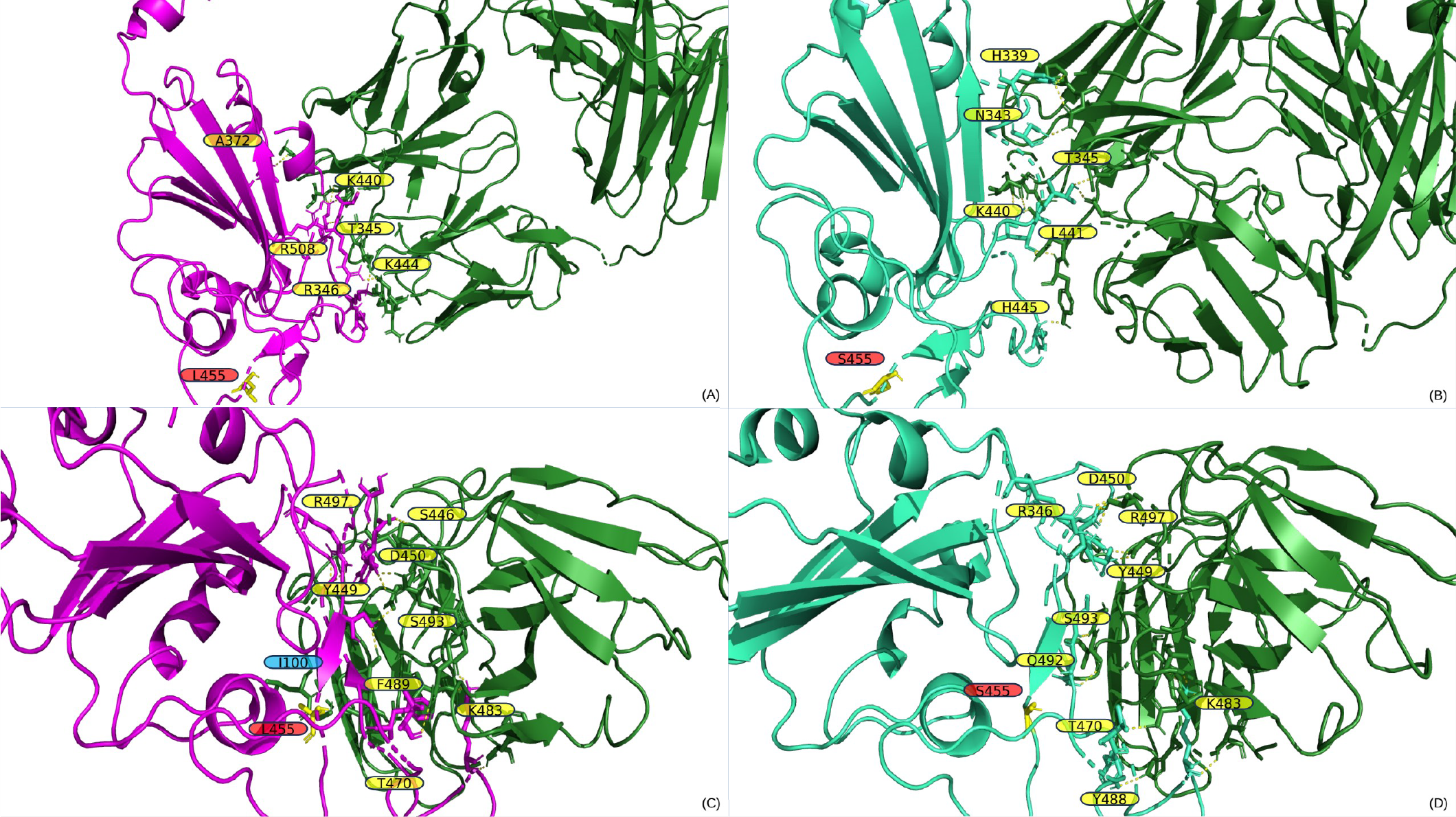
A) and B) display the output PDB files from HADDOCK of the docking jobs of antibody P2D9 to BA.2.86 and JN.1, respectively. C) and D) display the output PDB files from HADDOCK of the docking jobs of antibody 10-5B to BA.2.86 and JN.1, respectively. The antibody structure in each is shown on the right in green, and the RBD is shown on the left in magenta for BA.2.86 or greencyan for JN.1. Interacting RBD residues are labeled in yellow. Residues discussed in the results section on the antibody are labeled in blue. Important mutations referenced in the results section are highlighted in yellow and labeled in red.

Figure 4 shows the structural analysis of the interfacing residues between the RBDs of JN.1, BA.2.86, and XBB.1.5 with ACE2. When assessing Figure 4(A) and (B), one can notice that XBB.1.5 and JN.1 adopt a markedly different binding pose to each other, the angle of binding has approximately a 90-degree disparity. However, they do maintain similar interfacing residues. First, it is to note that at position 445, XBB.1.5 has a Proline residue while JN.1 has a Histidine residue. Proline offers enhanced rigidity to that of Histidine, potentially adding to the disparity in tertiary structure between XBB.1.5 and JN.1, affecting binding. Second, there is a deletion in the JN.1 RBD at position 483, where there is a Valine in position 483 in XBB.1.5. This Valine does not interface with ACE2. However, this deletion may lead to an alteration in tertiary structure that causes a variation in binding pose. Lastly, there is a Serine at position 490 in XBB.1.5, while there is a Phenylalanine in the corresponding position of 489 in JN.1. When assessing XBB.1.5, this Serine may form contacts in slight alterations of the docking pose with the Threonine27, Lysine31, and Glutamate35 in ACE2. These contacts could potentially increase stability. When assessing the corresponding structure of JN.1, the Phenylalanine is proximal to Histidine34 of ACE2. This proximity may increase pi-pi bonding. However, given the constraints of the Alpha Helix that Histidine34 is part of, there is limited flexibility, and steric hindrance between the two bulky side chains could cause detrimental effects on binding. When assessing Figure 4(C) and (D), one can notice a minimal difference in binding pose between BA.2.86 and JN.1. The only difference between the sequence of BA.2.86 and JN.1 is the residue at position 455. In JN.1, the residue at position 455 is a Leucine. This Leucine is close in proximity to Isoleucine100 and Phenylalanine72 in ACE2. This proximity could add to nonpolar interactions, increasing binding affinity. When assessing the Serine in position 455 in JN.1, it is positioned far away from the ACE2-RBD interface, leaving it unlikely to interact with any ACE2 residue to strengthen interaction significantly. There is a Lysine at position 68 in ACE2, which may form an interaction with the electronegative oxygen within Serine. However, Lysine68 is part of an alpha-helix of ACE2 and is angled away from Serine455 of the RBD, making the interaction less likely.

**Fig. 4.**
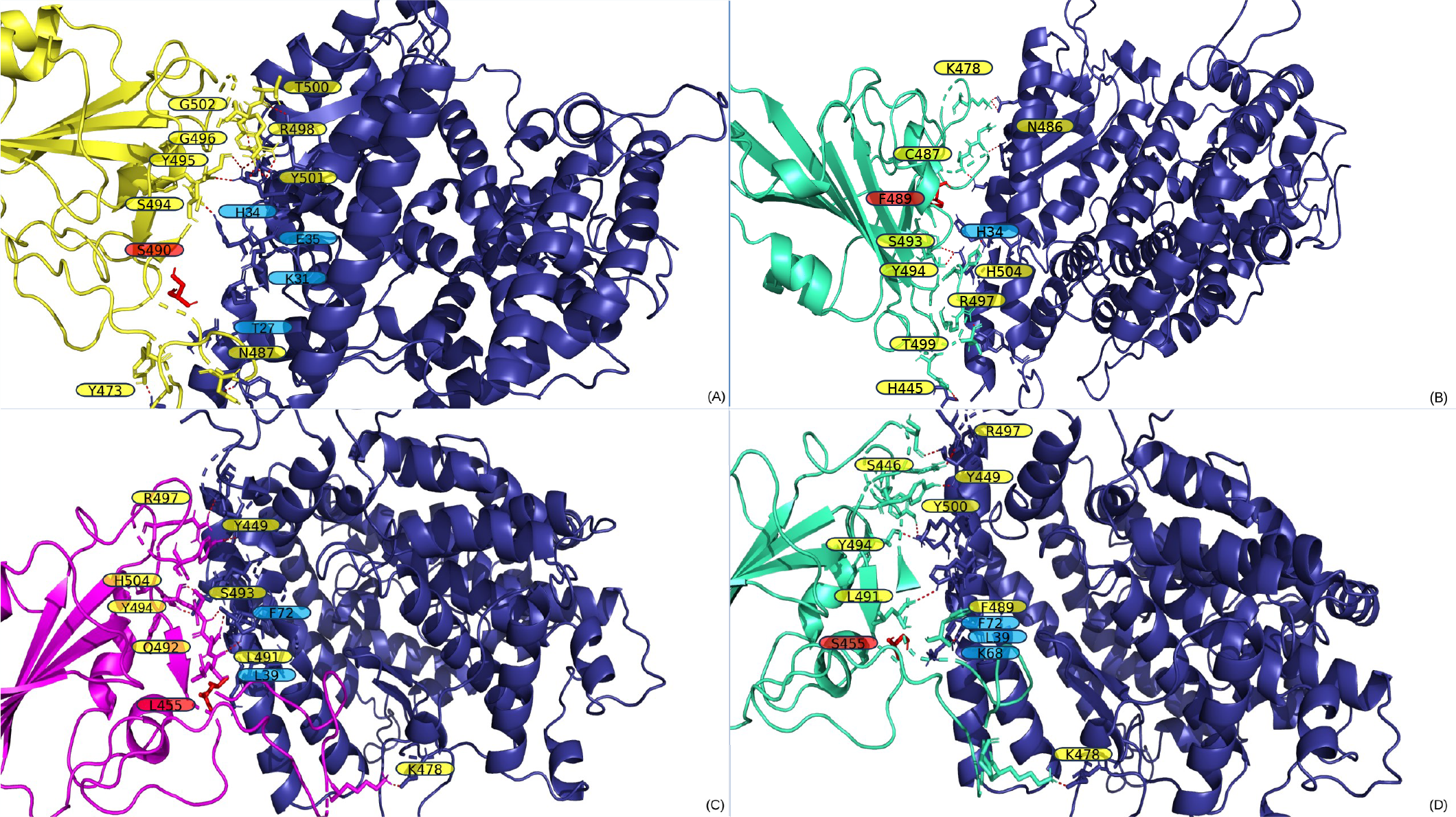
A) and B) display the output PDB files from HADDOCK of the docking jobs of the ACE2 structure from PDB 7XO9 and XBB.1.5 and JN.1, respectively. C) and D) display the output PDB files from HADDOCK of the docking jobs between the ACE2 structure from PDB 7WPB and BA.2.86 and JN.1, respectively. In each, the images show the ACE2 structure on the right in blue and the RBD on the left, colored in yellow for XBB.1.5, magenta for BA.2.86, and greencyan for JN.1. Interacting ACE2 residues, referenced in the results section, are labeled in blue. Important mutations referenced in the results section are labeled in red and highlighted in red.

### Broadly Neutralizing Antibody Performance

#### P2D9

The antibody P2D9, identified in Luo et al. 2023, was obtained from the memory B cells of individuals vaccinated with the BNT162b2-vaccine (31). These individuals also had breakthrough BA.1 infection (31). P2D9 was found to neutralize all tested variants of concern and omicron sublineages (31). Luo et al. tested neutralization capabilities against variants such as the WT, Alpha, Delta, BA.1, and BA.2 variants (31). Pseudovirus neutralization profiles were determined for P2D9. Luo et al. found on the BA.1, BA.2. and the WT variant, that P2D9 had IC50 values of 0.^0117, 0^.1381, and 0.0075 µg/mL, respectively (31). When assessing the values in Supplementary Table 3, one can notice similar values for Van der Waals energy and HADDOCK score between the three variants. However, the WT variant has a significantly lower Prodigy ΔG value, correlating with the empirical values. It is noted that BA.2 records a lower Prodigy ΔG value than the BA.1 Prodigy ΔG value. However, BA.1 Electrostatic Energy value is substantially lower than that of BA.2 potentially explaining the discrepancy between their IC50 values despite their Prodigy ΔG values. The WT variant has an Electrostatic Energy value between BA.1 and BA.2. The metrics in Supplementary Table 3 indicate P2D9 will perform similarly to previous Omicron variants on BA.2.86 and JN.1, maintaining its broadly neutralizing characteristic.

#### 6-2C

The antibody 6-2C, identified in Liu et al. 2023, was obtained from the memory B-cells of individuals vaccinated with the BBIBP-CorV inactivated vaccine (30). Specifically, Liu et al. obtained samples from patients who had robust humoral immune responses after the second dose of the BBIBP-CorV vaccine (30). When assessing Supplementary Table 4, one can notice that the WT variant has lower values for Van der Waals energy, HADDOCK score, and Prodigy ΔG relative to the BA.1 and BA.2 variant. Electrostatic energy is slightly higher between 6-2C and the WT variant. Overall, these metrics indicate that 6-2C has a higher binding affinity to the WT variant relative to BA.1 and BA.2 variants. When assessing empirical data reported in Liu et al. 2023, the minimum amount of antibody quantity to achieve 50% reduction in viral infectivity for the WT, BA.1, and BA.2 variants is 37, 1149, and 948 ng/mL respectively (30). One can note that 6-2C is far more potent on the WT variant than BA.1 and BA.2 variants. The empirical values correlate with the obtained docking metrics and post-docking metrics. When assessing the data for BA.2.86 and JN.1 in Supplementary Table 4, the metrics indicate that 6-2C will maintain neutralization capabilities for BA.2.86 and JN.1.

#### CV2.1169

The antibody CV2.1169, identified in Planchais et al. 2022, was obtained from the memory B-cells of individuals infected with the WT strain. Planchais et al. specifically filtered for serum samples from individuals with high seroneutralization for single B-cell antibody cloning. Plan-chais et al. found that the antibody CV2.1169 potently neutralized SARS-CoV-2 variants of concern. Planchais et al. concluded that CV2.1169 was a prime candidate for the prevention and treatment of COVID-19. When analyzing the Planchais et al. 2022 study, IC50 values for BA.1 and BA.2 are 850 and 756 pM, respectively, corresponding with the Prodigy ΔG values displayed in Supplementary Table 5, with BA.2 demonstrating a lower Prodigy ΔG, lower electrostatic energy score, and lower HADDOCK score. The data in Supplementary Table 5 indicates CV2.1169 will maintain its neutralizing capabilities as the various docking and affinity metrics for BA.2.86 and JN.1 are similar to BA.1 and BA.2.

## Discussion

The docking metrics we obtained tell a complex story when assessing them as a single component to form conclusions regarding the binding affinity of BA.2.86 and JN.1 to neutralizing antibodies and ACE2. Statistically significant Desolvation Energy differences are noted in the results. Desolvation Energy measures the energy change of protein atoms moving from interactions with a solvent to a non-solvent (56). Desolvation Energy can significantly impact binding affinity and is a component in the calculation of binding affinity (57). Taken alone, desolvation energy metrics may indicate lower or higher binding between two separate molecules within a solvent. However, we have the binding affinity metrics provided by PRODIGY, which do not indicate a statistically significant difference in binding affinity for neutralizing antibodies and ACE2 between JN.1 and BA.2.86 relative to previous Omicron variants.

Substantial changes in binding affinity are not observed between JN.1 and BA.2.86 to neutralizing antibodies and ACE2. BA.2.86 appears to have no significant evasion to RBD targeting antibodies and does not appear to bind with a significantly higher affinity to ACE2 than previous variants. Relative to BA.2.86, JN.1 does demonstrate a 3.9% increase in the median ΔG value to the RBD-targeting antibody arsenal in our study, indicating slightly increased antibody evasion. At the same time, JN.1 exhibits a lower ACE2 binding affinity than BA.2.86, with a 4.9% increase for the median ΔG value for JN.1 relative to BA.2.86.

### BA.2.86 Empirical Evidence Corresponds with Predictions

Studies have been published regarding BA.2.86 and JN.1. In one study titled, “Sensitivity of BA.2.86 to prevailing neutralizing antibody responses,” patients had blood sera collected before and after XBB.1.5 predominance (58). Sheward et al. found that blood samples collected before and after XBB.1.5 prevalence had moderately lower geometric mean neutralizing titers for BA.2.86 relative to XBB.1.5. Sheward et al. concluded that BA.2.86 did not appear to have the same level of relative immune escape as when the Omicron variant initially emerged in the shadow of the Delta variant in late 2021. The *in silico* results for BA.2.86 correspond with Sheward et al. 2023.

In an article published titled ‘SARS-CoV-2 Omicron sub-variant BA.2.86: limited potential for global spread’, Wang et al. review numerous studies regarding BA.2.86 including Sheward et al. 2023. Wang et al. analyze multiple studies that show that recent infection or vaccination increases neutralization of BA.2.86. Wang et al. generally conclude that BA.2.86 may not possess the transmissibility that previous Omicron strains possessed. Wang et al.’s analysis corresponds with the *in silico* results for BA.2.86. Wang et al. do note that their conclusion does not apply to the JN.1 variant as they did not examine it. Wang et al.’s review corresponds with the results of our study.

### Empirical Evidence Concerning Immune Evasion of JN.1 is Mixed

In an article titled ‘Humoral immune escape by current SARS-CoV-2 variants BA.2.86 and JN.1, December 2023’, Jeworowski et al. collected sera from a group of individuals who had received three to four vaccine doses in September 2023, a majority having prior breakthrough infection (20). These individuals were initially vaccinated using mRNA or vector-based vaccinations (20). Jeworowski et al. found that JN.1 did not exhibit a significant reduction in serum neutralization titers relative to BA.2.86 (20). It is to be noted that all of the individuals in the study reported a SARS-CoV-2 infection during the period of Omicron prevalence (20). Jeworowski et al. concluded that antibody evasion did not appear to cause the increased prevalence of JN.1. Jeworowski et al. 2024‘s results correspond with the results of our study.

In another study on JN.1 titled “Fast evolution of SARS-CoV-2 BA.2.86 to JN.1 under heavy immune pressure’, Yang et al. studied the humoral immune evasion and ACE2 binding affinity of JN.1 (17). Yang et al. collected blood samples from patients who were reinfected with XBB post BA.5 or BA.7 breakthrough infection (17). Yang et al. also collected samples from individuals who had received three doses of inactivated vaccines and subsequently contracted XBB (17). Yang et al. used a blood serum neutralization titer method to assess antibody neutralization. Yang et al. found that JN.1 had significantly higher antibody evasion than BA.2.86 but had a notably lower ACE2 binding affinity than BA.2.86 (17). Yang et al.‘s trends generally correspond with our results. However, the magnitude to which we observe antibody evasion is distinct (17). We do not note a significant increase, which Yang et al. note, in antibody evasion relative to previous strains, there is a noted decrease in ACE2 binding, as there is a reduction in binding affinity for JN.1 relative to BA.2.86.

### Evolution Outside of the RBD

We have correctly analyzed relative immune evasion for RBD targeting antibodies for the BA.1/B.1.1.529, XBB.1.5, and BA.2.86 variants with the *in silico* approach in the past (7, 21). When assessing the CDC Data Tracker, JN.1 accounts for approximately 96.4% of current variants detected (15). Our approach, in which we focus on RBD-antibody and RBD-ACE2 interaction, does not explain the relative predominance of JN.1 to BA.2.86.

Immune evasion and transmissibility are not exclusively dictated by mutations in the RBD region.

In the case of JN.1, several mutations of interest are present in nonstructural proteins that may play a role in immune evasion (13, 14, 60). For instance, mutations in nsp6 have been identified as capable of inhibiting the interferon type I (IFN-I) pathway (60). This inhibition leads to the absence of IFNA and IFNB production, crucial components in the defense against viral infections (61, 62).

In our tabulation of mutations 6, JN.1 contains a mutation, which is lacking in BA.2.86, in the proteins nsp3 and nsp6. JN.1 also contains a mutation, lacking in BA.2.86, within ORF7b, a gene whose function is not well studied. Nsp3 is associated with binding to host proteins and plays a role in viral replication (63).

This study, alongside our previous study involving XBB.1.5, demonstrates a decline in structural and electrostatic change within the RBD to evade vaccine, infection, and therapeutically derived antibodies (7). The predominant variants of the last two years maintain similar antibody evasion potential, marking a stark contrast to the structural and electrostatic divergence in the RBD that was introduced with the B.1.1.529 variant, which was highly evasive of existing antibodies, causing significant increases in hospitalizations and deaths (7, 15, 21). The results of our *in silico* analysis alongside traditional empirical analysis demonstrate that antibody evasion may not be the source of the relative increase in transmissibility demonstrated by JN.1 over BA.2.86 (20). Increased affinity to the ACE2 receptor is the direction of evolutionary pressure for the RBD of the Spike Protein. Antibody defenses derived from memory b-cells, via breakthrough infection and vaccination, have been studied to be increasingly more capable of binding the RBD of a broad amount of SARS-CoV-2 variants, which make their aggregate electrostatic and steric interactions increasingly similar to that of ACE2 (64). Therefore, since SARS-CoV-2 faces evolutionary pressure to evolve towards a highly conserved target, as ACE2 maintains conservation and a slow evolutionary rate, it is improbable that the RBD will evolve around increasingly strong antibody defenses without reducing ACE2 affinity, which would inherently reduce its transmissibility (65). We noted this very phenomenon with JN.1 relative to BA.2.86, as JN.1 has slightly increased antibody evasion, but it also has decreased ACE2 affinity. This substantial increase in predominance for JN.1 over BA.2.86, alongside our results and existing empirical results, implies that evolution outside of the RBD is enhancing relative SARS-CoV-2 variant transmissibility (20). Unfortunately, precise quantification of the impact of mutations outside the Spike protein contributing to JN.1’s increased immune escape compared to other variants remains challenging (17).

### Limitations and Advantages

We acknowledge that we cannot, as of yet, assess the interaction of multiple antibodies with the spike protein in neutralization interactions. However, a significant benefit to this approach is the ability to study the neutralization capabilities of individual antibodies, not the aggregate capabilities of antibodies prevalent in the blood. The antibody selection significantly differs from the antibodies studied in existing serological studies. The studied antibody array primarily consists of memory B-cell derived antibodies that are studied to be broadly neutralizing against different variants (30, 31, 37, 39, 42, 47). Memory B-cell-derived antibodies created in response to an antigen are not as easily observable when using blood serum neutralization titers. The *in silico* results support the empirical results that the selected antibody array, primarily consisting of broadly neutralizing antibodies, maintains its efficacy across different variants (30, 31, 37, 39, 42, 47). The presence of such broadly neutralizing antibodies also may indicate a level of prevalence of memory B-cells within vaccinated or in-fected individuals that differentiate to produce such antibodies (30, 37, 39, 42, 47, 66, 67). Serum antibody neutralization titers may be accurate in predicting the prevention of initial infection but are not as accurate in regards to the prediction of the prevention of serious disease. This is because serum antibody neutralization titers only account for antibodies prevalent in the blood at the time of collection. The secondary immune response, via the production of antibodies derived from memory B-cells, cannot easily be measured using this method. Neutralization titers, using blood samples, is a biased measurement method as the proportions of antibodies within the bloodstream will reflect specific neutralizing capabilities against recently introduced antigens from infection or vaccination. For example, in Yang et al. 2023, the authors specifically assess sera collected after XBB infection, biasing the proportions of antibodies within the blood that have neutralizing capabilities towards epitopes of XBB and structurally similar variants. When an immune response occurs to an introduced SARS-CoV-2 antigen in an individual with memory B-cells that can respond to that antigen, the proportions of neutralizing antibodies can and will change as the memory B-cells will differentiate to produce plasma cells to create neutralizing antibodies (66). In contrast to the limitation of a bias in the proportion of neutralizing antibodies provided in serum neutralization studies, this *in silico* method primarily uses antibodies that have been studied to be produced upon memory B-cell stimulation and neutralize SARS-CoV-2 variants, which may be more indicative of protection induced by prior infection or vaccination. A larger and broader study analyzing the presence of memory B-cells that differentiate to eventually produce broadly neutralizing antibodies is needed to support the notion that the general population can produce such broadly neutralizing and effective antibodies against current and future variants. In addition to the study of the natural production of broadly neutralizing antibodies, future work into developing therapeutic antibodies based upon the highly capable antibodies produced by the memory B-cells used in the study could lead to improved treatment outcomes.

## Conclusion

Our study indicates that ACE2 and antibody binding of the BA.2.86 and JN.1 variants is not profoundly different from previous variants. Moreover, it shows the ongoing efficacy of antibodies induced by various means in the global population to fight BA.2.86 and JN.1. Finally, the *in silico* results suggest that variation within the RBD has not led to the increased fitness JN.1 has demonstrated relative to BA.2.86. Mutations outside the RBD that may enhance fitness must be studied to enhance our ability to respond to the ongoing SARS-CoV-2 pandemic. Thus, it is important to approach genomic epidemiology with a functional perspective above and beyond the mission of sequencing novel variants.

## Data availability statement

All code, data, results, and additional analyses are openly available on GitHub at:

https://github.com/colbyford/SARS-CoV-2_BA.2.86_Spike-RBD_Predictions.

## Acknowledgements

The FP7 WeNMR (project # 261572), H2020 West-Life (project # 675858), the EOSC-hub (project # 777536) and the EGI-ACE (project # 101017567) European e-Infrastructure projects are acknowledged for the use of their web portals, which make use of the EGI infrastructure with the dedicated support of CESNET-MCC, INFN-LNL-2, NCG-INGRID-PT, TW-NCHC, CESGA, IFCA-LCG2, UA-BITP, TR-FC1-ULAKBIM, CSTCLOUD-EGI, IN2P3-CPPM, CIRMMP, SURFsara and NIKHEF, and the additional support of the national GRID Initiatives of Belgium, France, Italy, Germany, the Netherlands, Poland, Portugal, Spain, UK, Taiwan and the US Open Science Grid.

We acknowledge the Statens Serum Institut Bioinformatics and Microbial Genomics for the upload of the BA.2 genome sequence on GISAID.

We acknowledge the following entities at the University of North Carolina at Charlotte: Academic Affairs, The Office of Research, The Center for Computational Intelligence to Predict Health and Environmental Risks (CIPHER), The Department of Bioinformatics and Genomics, The College of Computing and Informatics, and the University Research Computing group. We acknowledge the support of the Belk Family.

## Supplementary Materials

**Table 3.** Individual docking metrics for the antibody P2D9 against the different variants used in the study.

| Variant | HADDOCK Score | Van der Waals energy | Electrostatic energy | Prodigy $\Delta G$ Kcal/mol |
| --- | --- | --- | --- | --- |
| WT | -85.5 | -72.1 | -248 | -13.3 |
| BA.1/B.1.1.529 | -88.7 | -65.5 | -285.8 | -11.5 |
| BA.2 | -88.2 | -70.1 | -204.9 | -12.2 |
| XBB.1.5 | -70.7 | -64.3 | -185.9 | -10.3 |
| BA.2.86 | -82.4 | -71.8 | -166.9 | -12.4 |
| JN.1 | -82.7 | -61.0 | -215.6 | -10.4 |

**Table 4.** Individual docking metrics for the antibody 6-2C against the different variants used in the study.

| Variant | HADDOCK score | Van der Waals energy | Electrostatic energy | Prodigy $\Delta G$ Kcal/mol |
| --- | --- | --- | --- | --- |
| WT | -106.4 | -86.1 | -205.8 | -14.2 |
| BA.1/B.1.1.529 | -73.3 | -54.9 | -229.4 | -9.9 |
| BA.2 | -90.7 | -68.8 | -216.4 | -13.4 |
| XBB.1.5 | -82.3 | -64.2 | -240.6 | -13.7 |
| BA.2.86 | -78.5 | -74.1 | -165.0 | -10.7 |
| JN.1 | -88.8 | -79.2 | -118.2 | -12.1 |

**Table 5.** Individual docking metrics for the antibody CV2.1169 against the different variants used in the study.

| Variant | HADDOCK Score | Van der Waals energy | Electrostatic energy | Prodigy $\Delta G$ Kcal/mol |
| --- | --- | --- | --- | --- |
| WT | -79.2 | -60.1 | -167.3 | -12.7 |
| BA.1/B.1.1.529 | -74.6 | -60.8 | -182.7 | -11.4 |
| BA.2 | -80.4 | -60.9 | -214 | -13.1 |
| XBB.1.5 | -81.8 | -65.4 | -175.7 | -13 |
| BA.2.86 | -94.7 | -67.7 | -251.9 | -12.7 |
| JN.1 | -112.8 | -80.6 | -230 | -14.1 |

**Table 6.**
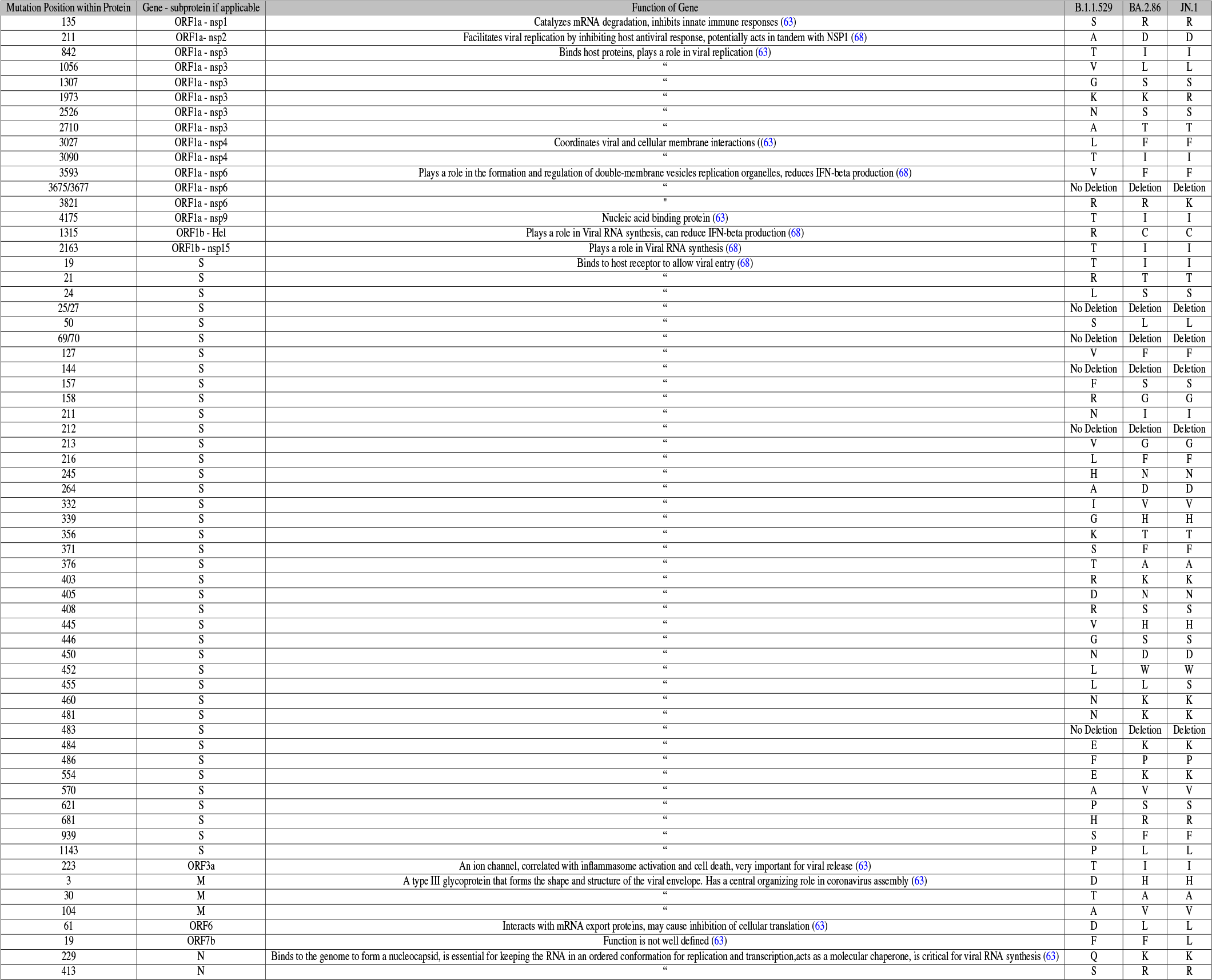
The table above displays the mutations for JN.1 and BA.2.86 relative to the B.1.1.529 variant. It shows the mutation location, the mutated amino acid present, the gene that the mutation is within, and the function of the gene. The mutation data is derived from the Outbreak.info lineage comparison tool and the UCSC Genome Browser (68, 69).

## Bibliography

1. UK Health Security Agency. Risk assessment for SARS-CoV-2 variant V-23AUG-01 (BA.2.86), Aug 2023. URL: https://www.gov.uk/government/publications/investigation-of-sars-cov-2-variants-of-concern-variant-risk-assessments/risk-assessment-for-sars-cov-2-variant-v-23aug-01-or-ba286; Accessed August 23, 2023.

2. Centers for Disease Control and Prevention. Risk Assessment Summary for SARS CoV-2 Sublineage BA.2.86., Aug 2023. URL: https://www.cdc.gov/respiratory-viruses/whats-new/covid-19-variant.html; Accessed November 1, 2023].

3. Suresh V. Kuchipudi. Why public health experts are concerned about BA.2.86, the latest COVID-19 variant, Sep 2023. URL: https://www.pbs.org/newshour/health/why-public-health-experts-are-concerned-about-ba-2-86-the-latest-covid-19-variant; Accessed: November 2, 2023.

4. Centers for Disease Control and Prevention. Update on SARS CoV-2 Variant BA.2.86, 2023. URL: https://www.cdc.gov/respiratory-viruses/whats-new/covid-19-variant-update-2023-08-30.html; Accessed August 10, 2023.

5. World Health Organization. Tracking SARS-CoV-2 variants, 2023. URL: https://www.who.int/activities/tracking-SARS-CoV-2-variants/; Accessed November 1, 2023].

6. Ewen Callaway. Coronavirus variant XBB.1.5 rises in the United States—is it a global threat? Nature, 613(7943):222–223, Jan 2023. doi: 10.1038/d41586-023-00014-3.

7. Colby T. Ford, Shirish Yasa, Denis Jacob Machado, Richard Allen White, and Daniel A. Janies. Predicting changes in neutralizing antibody activity for SARS-CoV-2 XBB.1.5 using in silico protein modeling. Frontiers in Virology, 3, 2023. ISSN 2673-818X. doi: 10.3389/fviro.2023.1172027.

8. Sijie Yang, Yuanling Yu, Fanchong Jian, Weiliang Song, Ayijiang Yisimayi, Xiaosu Chen, Yanli Xu, Peng Wang, Jing Wang, Lingling Yu, and et al. Antigenicity and infectivity characterisation of SARS-CoV-2 BA.2.86. The Lancet Infectious Diseases, 23(11), Sep 2023. doi: 10.1016/s1473-3099(23)00573-x.

9. Alice Park. Will the New COVID-19 Vaccine Work Against the BA.2.86 Variant?, 2023. URL: https://time.com/6308418/ba-2-86-covid-19-variant-vaccine/.

10. Christian A. Devaux and Jacques Fantini. ACE2 receptor polymorphism in humans and animals increases the risk of the emergence of SARS-CoV-2 variants during repeated intra- and inter-species host-switching of the virus. Front Microbiol, Jul 2023. doi: 10.3389/fmicb.2023.1199561.

11. Can Yue, Weiliang Song, Lei Wang, Fanchong Jian, Xiaosu Chen, Fei Gao, Zhongyang Shen, Youchun Wang, Xiangxi Wang, and Yunlong Cao. ACE2 binding and antibody evasion in enhanced transmissibility of XBB.1.5. The Lancet Infectious Diseases, 2023. doi: 10.1016/S1473-3099(23)00010-5.

12. Neil Bate, Christos G. Savva, Peter C. E. Moody, Edward A. Brown, Sian E. Evans, Jonathan K. Ball, John W. R. Schwabe, Julian E. Sale, and Nicholas P. J. Brindle. In vitro evolution predicts emerging SARS-CoV-2 mutations with high affinity for ACE2 and cross-species binding. Public Library of Science Pathogens, 2022. doi: 10.1371/journal.ppat.1010733.

13. Lianpan Dai, Tianyi Zheng, Kun Xu, Yuxuan Han, Lili Xu, Enqi Huang, Yaling An, Yingjie Cheng, Shihua Li, Mei Liu, et al. A universal design of betacoronavirus vaccines against covid-19, mers, and sars. Cell, 182(3):722–733, 2020.

14. Jingyun Yang, Wei Wang, Zimin Chen, Shuaiyao Lu, Fanli Yang, Zhenfei Bi, Linlin Bao, Fei Mo, Xue Li, Yong Huang, et al. A vaccine targeting the rbd of the s protein of sars-cov-2 induces protective immunity. Nature, 586(7830):572–577, 2020.

15. Centers of Disease Control and Prevention. COVID data tracker, Nov 2023. URL: https://covid.cdc.gov/covid-data-tracker/variant-proportions; Accessed February 17, 2024].

16. Xinling Wang, Lu Lu, and Shibo Jiang. SARS-CoV-2 evolution from the BA. 2.86 to JN. 1 variants: unexpected consequences. Trends in Immunology, 2024.

17. Sijie Yang, Yuanling Yu, Yanli Xu, Fanchong Jian, Weiliang Song, Ayijiang Yisimayi, Peng Wang, Jing Wang, Jingyi Liu, Lingling Yu, and et al. Fast evolution of SARS-COV-2 BA.2.86 to JN.1 under heavy immune pressure. The Lancet Infectious Diseases, 24(2), Dec 2023. doi: 10.1016/s1473-3099(23)00744-2.

18. Yu Kaku, Kaho Okumura, Miguel Padilla-Blanco, Yusuke Kosugi, Keiya Uriu, Alfredo A Hinay, Luo Chen, Arnon Plianchaisuk, Kouji Kobiyama, Ken J Ishii, et al. Virological characteristics of the SARS-CoV-2 JN. 1 variant. The Lancet Infectious Diseases, 2024.

19. Hang Fan, Si Qin, and Yujun Cui. Emergence and Characterization of the SARS-CoV-2 JN. 1 Variant: Global Prevalence and Implications for Public Health. Zoonoses, 4(1):994, 2024.

20. Lara M Jeworowski, Barbara Mühlemann, Felix Walper, Marie L Schmidt, Jenny Jansen, Andi Krumbholz, Etienne Simon-Lorière, Terry C Jones, Victor M Corman, and Christian Drosten. “Humoral immune escape by current SARS-CoV-2 variants BA.2.86 and JN.1, December 2023”. Eurosurveillance, 29(2):2300740, 2024. doi: 10.2807/1560-7917.ES.2024.29.2.2300740.

21. Colby T. Ford, Denis Jacob Machado, and Daniel A. Janies. Predictions of the SARS-CoV-2 Omicron variant (B.1.1.529) spike protein receptor-binding domain structure and neutralizing antibody interactions. Frontiers in Virology, 2, Feb 2022. doi: 10.3389/fviro.2022.830202.

22. Shruti Khare, Céline Gurry, Lucas Freitas, Mark B Schultz, Gunter Bach, Amadou Diallo, Nancy Akite, Joses Ho, Raphael TC Lee, Winston Yeo, GISAID Core Curation Team, and Sebastian Maurer-Stroh. GISAID’s role in pandemic response. China CDC Weekly, 3:1049, 2021. ISSN 2096-7071. doi: 10.46234/ccdcw2021.255.

23. Denis Jacob Machado, Rachel Scott, Sayal Guirales, and Daniel A Janies. Fundamental evolution of all Orthocoronavirinae including three deadly lineages descendent from Chiroptera-hosted coronaviruses: SARS-CoV, MERS-CoV and SARS-CoV-2. Cladistics, 37(5):461–488, 2021.

24. Shruti Khare, Céline Gurry, Lucas Freitas, Mark B Schultz, Gunter Bach, Amadou Diallo, Nancy Akite, Joses Ho, Raphael Tc Lee, Winston Yeo, Gisaid Core Curation Team, and Sebastian Maurer-Stroh. Perspectives: GISAID’s role in pandemic response. China CDC Weekly., 3(49):1049–1051, 2021. doi: 10.46234%2Fccdcw2021.255.

25. John Jumper, Richard Evans, Alexander Pritzel, Tim Green, Michael Figurnov, Olaf Ronneberger, Kathryn Tunyasuvunakool, Russ Bates, Augustin žídek, Anna Potapenko, et al. Highly accurate protein structure prediction with AlphaFold. Nature, 596(7873):583–589, 2021. doi: 10.1038/s41586-021-03819-2.

26. Milot Mirdita, Konstantin Schütze, Yoshitaka Moriwaki, Lim Heo, Sergey Ovchinnikov, and Martin Steinegger. ColabFold: making protein folding accessible to all. Nature Methods, 19 (6):679–682, Jun 2022. ISSN 1548-7105. doi: 10.1038/s41592-022-01488-1.

27. David A. Case, Hasan Metin Aktulga, Kellon Belfon, David S. Cerutti, G. Andrés Cisneros, Vinícius Wilian D. Cruzeiro, Negin Forouzesh, Timothy J. Giese, Andreas W. Götz, Holger Gohlke, Saeed Izadi, Koushik Kasavajhala, Mehmet C. Kaymak, Edward King, Tom Kurtzman, Tai-Sung Lee, Pengfei Li, Jian Liu, Tyler Luchko, Ray Luo, Madushanka Manathunga, Matias R. Machado, Hai Minh Nguyen, Kurt A. O’Hearn, Alexey V. Onufriev, Feng Pan, Sergio Pantano, Ruxi Qi, Ali Rahnamoun, Ali Risheh, Stephan Schott-Verdugo, Akhil Shajan, Jason Swails, Junmei Wang, Haixin Wei, Xiongwu Wu, Yongxian Wu, Shi Zhang, Shiji Zhao, Qiang Zhu, Thomas E. Cheatham III, Daniel R. Roe, Adrian Roitberg, Carlos Simmerling, Darrin M. York, Maria C. Nagan, and Kenneth M. Merz Jr. AmberTools. Journal of Chemical Information and Modeling, 63(20):6183–6191, 2023. doi: 10.1021/acs.jcim.3c01153.

28. Helen M. Berman, John Westbrook, Zukang Feng, Gary Gilliland, T. N. Bhat, Helge Weissig, Ilya N. Shindyalov, and Philip E. Bourne. The Protein Data Bank. Nucleic Acids Research, 28(1):235–242, 2000. ISSN 0305-1048. doi: 10.1093/nar/28.1.235.

29. Jun Lan, Jiwan Ge, Jinfang Yu, Sisi Shan, Huan Zhou, Shilong Fan, Qi Zhang, Xuanling Shi, Qisheng Wang, Linqi Zhang, and Xinquan Wang. Structure of the SARS-CoV-2 spike receptor-binding domain bound to the ACE2 receptor. Nature, 581(7807):215–220, May 2020. ISSN 1476-4687. doi: 10.1038/s41586-020-2180-5.

30. Yubin Liu, Ziyi Wang, Xinyu Zhuang, Shengnan Zhang, Zhicheng Chen, Yan Zou, Jie Sheng, Tianpeng Li, Wanbo Tai, Jinfang Yu, Yanqun Wang, Zhaoyong Zhang, Yunfeng Chen, Liangqin Tong, Xi Yu, Linjuan Wu, Dong Chen, Renli Zhang, Ningyi Jin, Weijun Shen, Jincun Zhao, Mingyao Tian, Xinquan Wang, and Gong Cheng. Inactivated vaccine-elicited potent antibodies can broadly neutralize SARS-CoV-2 circulating variants. Nature Communications, 14(1):2179, Apr 2023. ISSN 2041-1723. doi: 10.1038/s41467-023-37926-7.

31. Mengxiao Luo, Biao Zhou, Eswar R. Reddem, Bingjie Tang, Bohao Chen, Runhong Zhou, Hang Liu, Lihong Liu, Phinikoula S. Katsamba, Ka-Kit Au, Hiu-On Man, Kelvin Kai-Wang To, Kwok-Yung Yuen, Lawrence Shapiro, Shangyu Dang, David D. Ho, and Zhiwei Chen. Structural insights into broadly neutralizing antibodies elicited by hybrid immunity against SARS-CoV-2. Emerging Microbes & Infections, 12(1):2146538, Dec 2023. doi: 10.1080/22221751.2022.2146538.

32. Christopher O. Barnes, Claudia A. Jette, Morgan E. Abernathy, Kim-Marie A. Dam, Shannon R. Esswein, Harry B. Gristick, Andrey G. Malyutin, Naima G. Sharaf, Kathryn E. Huey-Tubman, Yu E. Lee, Davide F. Robbiani, Michel C. Nussenzweig, Anthony P. West, and Pamela J. Bjorkman. SARS-CoV-2 neutralizing antibody structures inform therapeutic strategies. Nature, 588(7839):682–687, Dec 2020. ISSN 1476-4687. doi: 10.1038/s41586-020-2852-1.

33. Christopher O Barnes, Anthony P West, Jr, Kathryn E Huey-Tubman, Magnus A G Hoffmann, Naima G Sharaf, Pauline R Hoffman, Nicholas Koranda, Harry B Gristick, Christian Gaebler, Frauke Muecksch, Julio C Cetrulo Lorenzi, Shlomo Finkin, Thomas Hägglöf, Arlene Hurley, Katrina G Millard, Yiska Weisblum, Fabian Schmidt, Theodora Hatziioannou, Paul D Bieniasz, Marina Caskey, Davide F Robbiani, Michel C Nussenzweig, and Pamela J Bjorkman. Structures of human antibodies bound to SARS-CoV-2 spike reveal common epitopes and recurrent features of antibodies. Cell, 182(4):828–842.e16, 2020. doi: 10.1016/j.cell.2020.06.025.

34. Donald J. Benton, Antoni G. Wrobel, Pengqi Xu, Chloë Roustan, Stephen R. Martin, Peter B. Rosenthal, John J. Skehel, and Steven J. Gamblin. Receptor binding and priming of the spike protein of SARS-CoV-2 for membrane fusion. Nature, 588(7837):327–330, 2020. ISSN 1476-4687. doi: 10.1038/s41586-020-2772-0.

35. Zhennan Zhao, Yufeng Xie, Bin Bai, Chunliang Luo, Jingya Zhou, Weiwei Li, Yumin Meng, Linjie Li, Dedong Li, Xiaomei Li, Xiaoxiong Li, Xiaoyun Wang, Junqing Sun, Zepeng Xu, Yeping Sun, Wei Zhang, Zheng Fan, Xin Zhao, Linhuan Wu, Juncai Ma, Odel Y. Li, Guijun Shang, Yan Chai, Kefang Liu, Peiyi Wang, George F. Gao, and Jianxun Qi. Structural basis for receptor binding and broader interspecies receptor recognition of currently circulating Omicron sub-variants. Nature Communications, 14(1):4405, 2023. ISSN 2041-1723. doi: 10.1038/s41467-023-39942-z.

36. Wei Zhang, Ke Shi, Qibin Geng, Morgan Herbst, Michael Wang, Linfen Huang, Fan Bu, Bin Liu, Hideki Aihara, and Fang Li. Structural evolution of SARS-CoV-2 omicron in human receptor recognition. Journal of Virology, 97(8):e00822–23, 2023. doi: 10.1128/jvi.00822-23.

37. Kang Wang, Zijing Jia, Linilin Bao, Lei Wang, Lei Cao, Hang Chi, Yaling Hu, Qianqian Li, Yunjiao Zhou, Yinan Jiang, Qianhui Zhu, Yongqiang Deng, Pan Liu, Nan Wang, Lin Wang, Min Liu, Yurong Li, Boling Zhu, Kaiyue Fan, Wangjun Fu, Peng Yang, Xinran Pei, Zhen Cui, Lili Qin, Pingju Ge, Jiajing Wu, Shuo Liu, Yiding Chen, Weijin Huang, Qiao Wang, Cheng-Feng Qin, Youchun Wang, Chuan Qin, and Xiangxi Wang. Memory B cell repertoire from triple vaccinees against diverse SARS-CoV-2 variants. Nature, 603(7903):919–925, Mar 2022. ISSN 1476-4687. doi: 10.1038/s41586-022-04466-x.

38. Cong Xu, Yanxing Wang, Caixuan Liu, Chao Zhang, Wenyu Han, Xiaoyu Hong, Yifan Wang, Qin Hong, Shutian Wang, Qiaoyu Zhao, Yalei Wang, Yong Yang, Kaijian Chen, Wei Zheng, Liangliang Kong, Fangfang Wang, Qinyu Zuo, Zhong Huang, and Yao Cong. Conformational dynamics of SARS-CoV-2 trimeric spike glycoprotein in complex with receptor ACE2 revealed by cryo-EM. Science Advances, 7(1):eabe5575, 2021. doi: 10.1126/sciadv.abe5575.

39. Cyril Planchais, Ignacio Fernández, Timothée Bruel, Guilherme Dias de Melo, Matthieu Prot, Maxime Beretta, Pablo Guardado-Calvo, Jérémy Dufloo, Luis M Molinos-Albert, Marija Backovic, et al. Potent human broadly SARS-CoV-2–neutralizing IgA and IgG antibodies effective against Omicron BA. 1 and BA. 2. Journal of Experimental Medicine, 219(7), 2022.

40. Kathryn Westendorf, Stefanie žentelis, Lingshu Wang, Denisa Foster, Peter Vaillancourt, Matthew Wiggin, Erica Lovett, Robin van der Lee, Jörg Hendle, Anna Pustilnik, et al. LY-CoV1404 (bebtelovimab) potently neutralizes SARS-CoV-2 variants. Cell reports, 39(7), 2022.

41. Bryan E Jones, Patricia L Brown-Augsburger, Kizzmekia S Corbett, Kathryn Westendorf, Julian Davies, Thomas P Cujec, Christopher M Wiethoff, Jamie L Blackbourne, Beverly A Heinz, Denisa Foster, et al. The neutralizing antibody, LY-CoV555, protects against SARS-CoV-2 infection in nonhuman primates. Science translational medicine, 13(593):eabf1906, 2021.

42. Rungtiwa Nutalai, Daming Zhou, Aekkachai Tuekprakhon, Helen M. Ginn, Piyada Supasa, Chang Liu, Jiandong Huo, Alexander J. Mentzer, Helen M.E. Duyvesteyn, Aiste Dijokaite-Guraliuc, and et al. Potent cross-reactive antibodies following omicron breakthrough in vaccinees. Cell, 185(12), May 2022. doi: 10.1016/j.cell.2022.05.014.

43. Pengcheng Han, Linjie Li, Sheng Liu, Qisheng Wang, D. Zhang, Zepeng Xu, Pu Han, Xiaomei Li, Qi Peng, Chao Su, et al. Receptor binding and complex structures of human ACE2 to spike RBD from omicron and delta SARS-CoV-2. Cell, 185(4):630–640, 2022.

44. Wanchao Yin, Youwei Xu, Peiyu Xu, Xiaodan Cao, Canrong Wu, Chunyin Gu, Xinheng He, Xiaoxi Wang, Sijie Huang, Qingning Yuan, et al. Structures of the Omicron spike trimer with ACE2 and an anti-Omicron antibody. Science, 375(6584):1048–1053, 2022.

45. James W Saville, Dhiraj Mannar, Xing Zhu, Alison M Berezuk, Spencer Cholak, Katharine S Tuttle, Faezeh Vahdatihassani, and Sriram Subramaniam. Structural analysis of receptor engagement and antigenic drift within the BA. 2 spike protein. Cell Reports, 42(1), 2023.

46. Youwei Xu, Canrong Wu, Xiaodan Cao, Chunyin Gu, Heng Liu, Mengting Jiang, Xiaoxi Wang, Qingning Yuan, Kai Wu, Jia Liu, et al. Structural and biochemical mechanism for increased infectivity and immune evasion of Omicron BA. 2 variant compared to BA. 1 and their possible mouse origins. Cell Research, 32(7):609–620, 2022.

47. Wanwisa Dejnirattisai, Daming Zhou, Helen M Ginn, Helen ME Duyvesteyn, Piyada Supasa, James Brett Case, Yuguang Zhao, Thomas S Walter, Alexander J Mentzer, Chang Liu, et al. The antigenic anatomy of SARS-CoV-2 receptor binding domain. Cell. doi: 10.1016/j.cell.2021.02.032.

48. Cyril Dominguez, Rolf Boelens, and Alexandre M. J. J. Bonvin. HADDOCK: a protein-protein docking approach based on biochemical or biophysical information. Journal of the American Chemical Society, 125(7):1731–1737, Feb 2003. ISSN 0002-7863. doi: 10.1021/ja026939x.

49. G.C.P. van Zundert, J.P.G.L.M. Rodrigues, M. Trellet, C. Schmitz, P.L. Kastritis, E. Karaca, A.S.J. Melquiond, M. van Dijk, S.J. de Vries, and A.M.J.J. Bonvin. The HADDOCK2.2 web server: user-friendly integrative modeling of biomolecular complexes. Journal of Molecular Biology, 428(4):720–725, 2016. doi: 10.1016/j.jmb.2015.09.014. Computation Resources for Molecular Biology.

50. Anna Vangone and Alexandre MJJ Bonvin. Contacts-based prediction of binding affinity in protein–protein complexes. eLife, 4:e07454, 2015. doi: 10.7554/eLife.07454.

51. Li C. Xue, João Pglm Rodrigues, Panagiotis L. Kastritis, Alexandre Mjj Bonvin, and Anna Vangone. PRODIGY: a web server for predicting the binding affinity of protein–protein complexes. Bioinformatics, 32(23):3676–3678, 08 2016. ISSN 1367-4803. doi: 10.1093/bioinformatics/btw514.

52. R Core Team. R: A Language and Environment for Statistical Computing. R Foundation for Statistical Computing, Vienna, Austria, 2022.

53. William H. Kruskal and W. Allen Wallis. Use of ranks in one-criterion variance analysis. Journal of the American Statistical Association, 47(260):583–621, 1952. doi: 10.1080/01621459.1952.10483441.

54. Frank Wilcoxon. Individual comparisons by ranking methods. Biometrics Bulletin, 1(6):80, 1945. doi: 10.2307/3001968.

55. Schrödinger, LLC. The PyMOL molecular graphics system, version 1.8. November 2015.

56. Hao Li, Yumeng Yan, Xuejun Zhao, and Sheng-You Huang. Inclusion of desolvation energy into protein–protein docking through atomic contact potentials. Journal of Chemical Information and Modeling, 62(3):740–750, 2022.

57. Brian K Shoichet, Andrew R Leach, and Irwin D Kuntz. Ligand solvation in molecular docking. Proteins: Structure, Function, and Bioinformatics, 34(1):4–16, 1999.

58. Daniel J Sheward, Yiqiu Yang, Michelle Westerberg, Sofia Öling, Sandra Muschiol, Kenta Sato, Thomas P Peacock, Gunilla B Karlsson Hedestam, Jan Albert, and Ben Murrell. Sensitivity of the sars-cov-2 ba. 2.86 variant to prevailing neutralising antibody responses. The Lancet Infectious Diseases, 23(11):e462–e463, 2023.

59. Xinling Wang, Lu Lu, and Shibo Jiang. SARS-CoV-2 Omicron subvariant BA. 2.86: limited potential for global spread. Signal Transduction and Targeted Therapy, 8(1):439, 2023.

60. Hongjie Xia, Zengguo Cao, Xuping Xie, Xianwen Zhang, John Yun-Chung Chen, Hualei Wang, Vineet D Menachery, Ricardo Rajsbaum, and Pei-Yong Shi. Evasion of type i interferon by sars-cov-2. Cell reports, 33(1), 2020.

61. Yoji Tsugawa, Hiroki Kato, Takashi Fujita, Kunitada Shimotohno, and Makoto Hijikata. Critical role of interferon-αconstitutively produced in human hepatocytes in response to rna virus infection. PLoS One, 9(2):e89869, 2014.

62. Sayuri Sakuragi, Huanan Liao, Kodai Yajima, Shigeyoshi Fujiwara, and Hiroyuki Nakamura. Rubella virus triggers type i interferon antiviral response in cultured human neural cells: involvement in the control of viral gene expression and infectious progeny production. International Journal of Molecular Sciences, 23(17):9799, 2022.

63. Chongzhi Bai, Qiming Zhong, and George Fu Gao. Overview of SARS-CoV-2 genome-encoded proteins. Science China Life Sciences, 65(2):280–294, 2022.

64. Shiru Chen, Fei Guan, Fabio Candotti, Kamel Benlagha, Niels Olsen Saraiva Camara, Andres A Herrada, Louisa K James, Jiahui Lei, Heather Miller, Masato Kubo, et al. The role of B cells in COVID-19 infection and vaccination. Frontiers in immunology, 13:988536, 2022.

65. Sally Badawi and Bassam R Ali. ACE2 Nascence, trafficking, and SARS-CoV-2 pathogenesis: the saga continues. Human genomics, 15(1):8, 2021.

66. Isaak Quast and David Tarlinton. B cell memory: understanding COVID-19. Immunity, Jan 2021. doi: 10.1016/j.immuni.2021.01.014.

67. Zeli Zhang, Jose Mateus, Camila H. Coelho, Jennifer M. Dan, Carolyn Rydyznski Moderbacher, Rosa Isela Gálvez, Fernanda H. Cortes, Alba Grifoni, Alison Tarke, James Chang, E. Alexandar Escarrega, Christina Kim, Benjamin Goodwin, Nathaniel I. Bloom, April Frazier, Daniela Weiskopf, Alessandro Sette, and Shane Crotty. Humoral and cellular immune memory to four COVID-19 vaccines. Cell, 185(14):2434–2451.e17, July 2022. doi: 10.1016/j.cell.2022.05.022.

68. W James Kent, Charles W Sugnet, Terrence S Furey, Krishna M Roskin, Tom H Pringle, Alan M Zahler, and David Haussler. The human genome browser at UCSC. Genome research, 12(6):996–1006, 2002. ‘Date accessed: 04/04/2024’.

69. Karthik Gangavarapu, Alaa Abdel Latif, Julia L Mullen, Manar Alkuzweny, Emory Hufbauer, Ginger Tsueng, Emily Haag, Mark Zeller, Christine M Aceves, Karina Zaiets, et al. Outbreak. info genomic reports: scalable and dynamic surveillance of SARS-CoV-2 variants and mutations. Nature methods, 20(4):512–522, 2023. “Accessed on 04/04/2024”.

